# Prebiotic Gas Flow Environment Enables Isothermal Nucleic Acid Replication

**DOI:** 10.1101/2024.06.13.598889

**Authors:** Philipp Schwintek, Emre Eren, Christof Mast, Dieter Braun

**Affiliations:** Systems Biophysics, Physics department, Center for NanoScience, Ludwig-Maximilians-Universität München, Amalienstraße 54, 80799 Munich, Germany

**Keywords:** molecular evolution, DNA, nucleic acids, origins of life, non-equilibrium, isothermal, replication, gas flux

## Abstract

Nucleic acid replication is a central process at the origin of life. On early Earth, replication is challenged by the dilution of molecular building blocks and the difficulty of separating daughter from parent strands, a necessity for exponential replication. While thermal gradient systems have been shown to address these problems, elevated temperatures lead to degradation. Also, compared to constant temperature environments, such systems are rare. The isothermal system studied here models an abundant geological environment of the prebiotic Earth, in which water is continuously evaporated at the point of contact with the gas flows, inducing up-concentration and circular flow patterns at the gas-water interface through momentum transfer. We show experimentally that this setting drives a 30-fold accumulation of nucleic acids and their periodic separation by a 3-fold reduction in salt and product concentration. Fluid dynamic simulations agree with observations from tracking fluorescent beads. In this isothermal system, we were able to drive exponential DNA replication with Taq polymerase. The results provide a model for a ubiquitous non-equilibrium system to host early Darwinian molecular evolution at constant temperature.

## Introduction

The emergence of life on Earth is still an unsolved puzzle to contemporary research. It is estimated that this event dates back approximately 3.7 - 4.5 billion years, with fossil carbon isotope signatures being the oldest evidence for life around 3.7 billion years ago [1, 2]. In order to reconstruct how early molecular life began before this time, it is crucial to identify and understand plausible geological environments, which support early prebiotic reaction networks that could have lead to the life we know today [3].

The common theory is that the Darwinian evolution of informational polymers was at the core of the origin of life [3]. Among these, nucleic acids, like RNA, stand out for their capability to both store genetic information and catalyze their own replication through transient formation of double-stranded helices [4]. These abilities allow them to mutate and evolve, enabling them to adapt to diverse environments and eventually encode, build and utilize proteins as the catalysts used in modern life.

Dilution, however, poses a significant obstacle, since such prebiotic reactions require sufficiently high concentrations of their reagents to work [5]. Large reservoirs, such as the ocean, cannot compensate for diffusion, because they lack local sources of energy to drive reaction pathways out of equilibrium [6]. The resulting homogeneity renders these environments unlikely to have harbored early molecular life [7].

Local physical non-equilibria, however, have shown the ability to up-concentrate molecules, such as nucleic acids, in a variety of different geological settings [8]. Examples range from thermal gradients in rock pores, local evaporation, re-hydration cycles of warm ponds, adsorption to mineral surfaces, heated gas bubbles in porous rocks, foams, or the eutectic phase in freeze-thaw cycles [9–17].

However, the accumulation of salts and molecules comes at a cost. Single-stranded nucleic acids replicate into double-stranded forms. These strands must separate again to complete a full replication cycle. But strand separation becomes increasingly difficult after accumulation, because the melting temperature of oligonucleotides is strongly dependent on the local salt concentration [18]. Despite high Mg^2+^ concentrations being required for replication and catalytic activity [19], they can elevate the melting temperature of nucleic acid duplex structures to levels surpassing even the boiling point of water [20]. Oligonucleotides readily hydrolyze into nucleotide fragments under these conditions, rendering high temperature spikes as a primary strand separation mechanism more detrimental than beneficial [21].

Therefore, other mechanisms are required at the origin of life to separate nucleic acid strands with minimal thermal stress, and at best combined with an environment where supplied biomolecules are accumulated from the environment and trapped for long periods of time. Examples have used pH oscillations to drive nucleic acid strand separation, which can be caused either by differential thermophoresis of ionic species or by periodic freeze-thaw cycles [22–24]. Also, dew droplet cycles in a rock pore subjected to a temperature gradient can periodically melt strands by transiently lowering the salt concentration [25, 26]. Heated gas-water interfaces were also shown to promote many prebiotic synthesis reactions [14, 27, 28]. The above scenarios require temperature gradients or thermal cycling. This creates degradation stress for nucleic acids and limits the scenarios to geological settings with a thermal gradient. Here, we investigated a simple and ubiquitous scenario in which a water flux through a rock pore was dried by a gas flux at constant temperature (Fig. 1). This can be found in the vicinity of underwater degassing events, where gases percolate through rocks to reach the surface, or in porous rocks at the surface exposed to atmospheric winds [29, 30]. Such a setting would be very common on volcanic islands on early Earth which also offered the necessary dry conditions for RNA synthesis[31].

**Figure 1.**
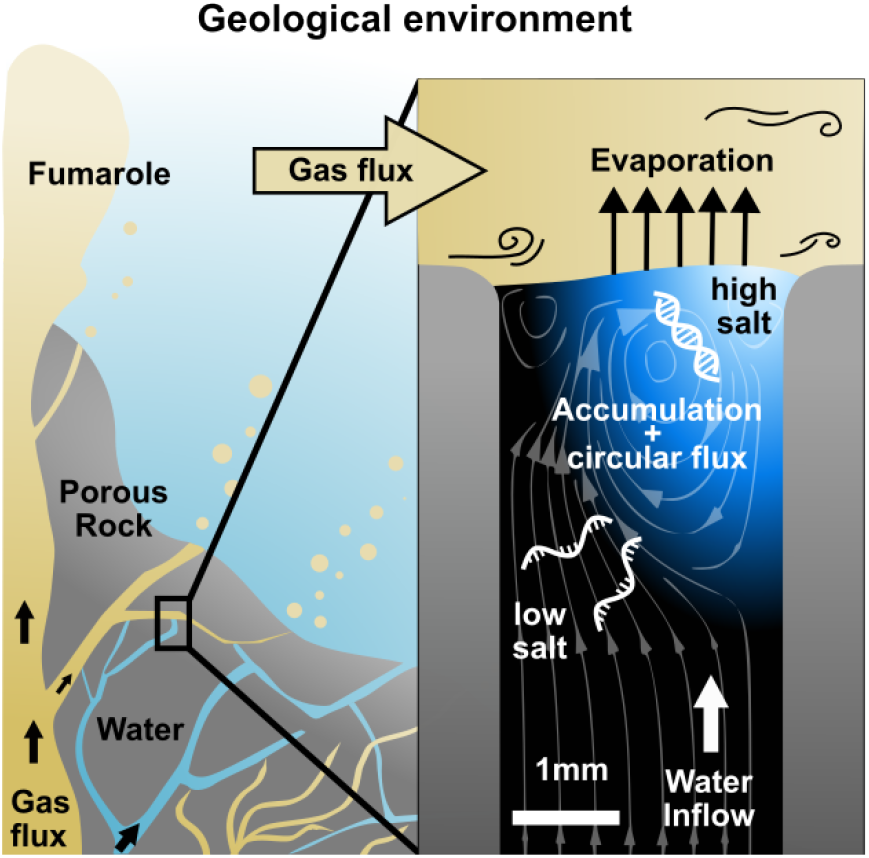
Replication at the gas-water interface. We considered a geological scenario in which water, containing biomolecules, is evaporated by a gas flow at the scale of millimeters. In volcanic porous rock, many of such settings can be imagined. The gas flow induces convective water currents and causes it to evaporate. Dissolved nucleic acids and salts accumulate at the gas-water interface due to the interfacial currents, even if the influx from below is pure water. Through the induced vortex, nucleic acids pass through different concentrations of salt, promoting strand separation and allowing them to replicate exponentially. Our experiments replicate this environment on the microscale, subjecting a defined sample volume to a continuous influx of pure water with an airflux brushing across.

We created an experimental model of such an evaporation pore, shown in Fig. 1, and studied how combined gas and water fluxes can lead to early replication of nucleic acids. We first analyzed accumulation flow speeds at the interface in Fig. 2, then monitored cyclic strand separation dynamics in Fig. 3, and finally showed how both drive DNA-based replication under isothermal conditions in Fig. 4.

**Figure 2.**
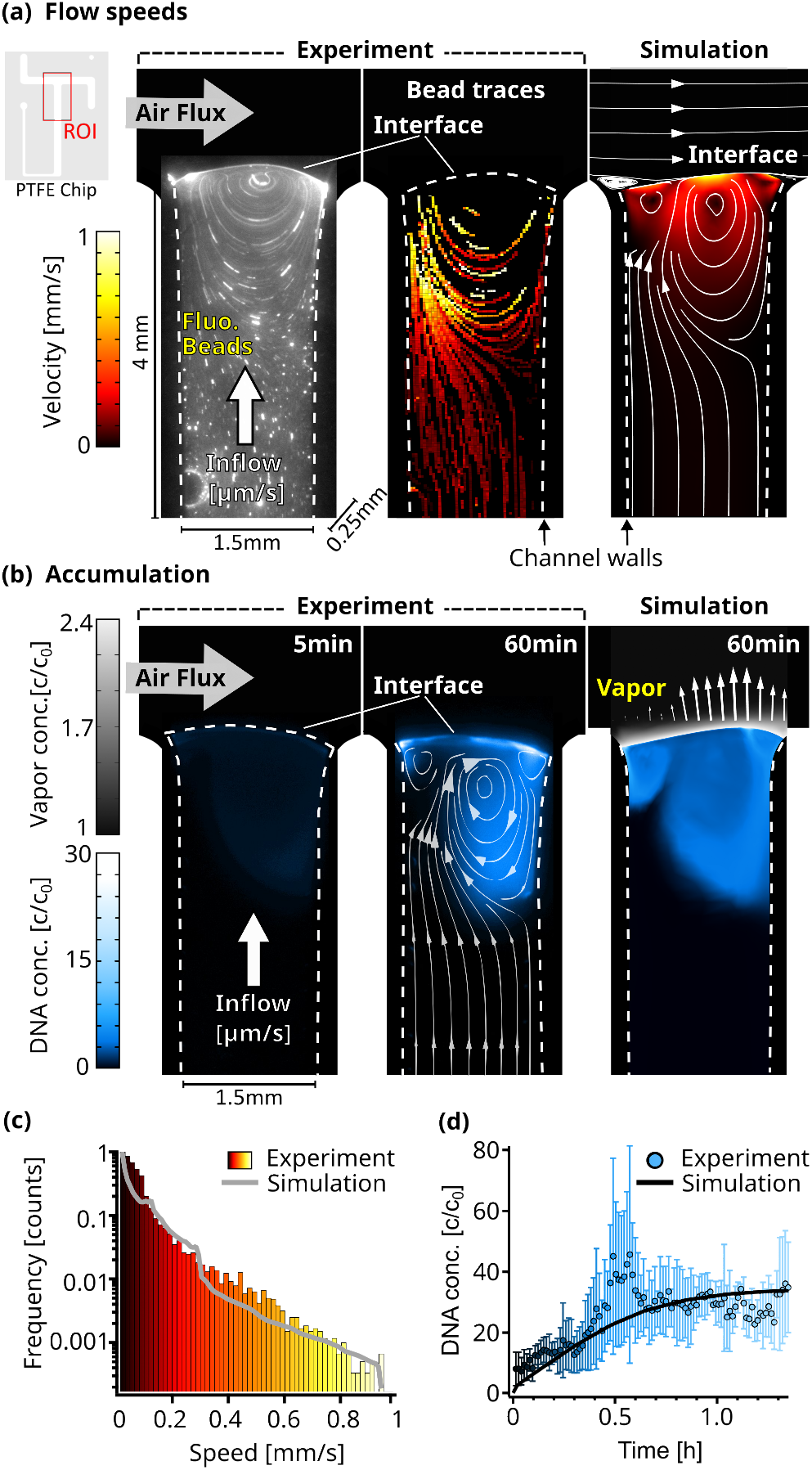
Flow and accumulation dynamics. **(a)** Imaging of fluorescent beads (0.5 μm) reveals a flow vortex right below the air-water interface, induced by the air flux across the interface (left panel). The bead movements were traced (middle panel) and the measured velocities were confirmed by a detailed finite element simulation (right panel). The PTFE chip cutout in the top left corner shows the ROI used for the micrographs. The color scale is equal for both simulation and experiment and Channel dimensions are 4 × 1.5 × 0.25 mm as indicated. Dotted lines visualize the location of the channel walls. **(b)** The accumulation of fluorescently labeled 63mer DNA was imaged and confirmed our understanding of the environment based on a diffusion model. Concentration reaches up to 30 times relative to the start c0. The accumulation profile of the experiment (middle panel) and simulation (right panel) match well, showcased by overlaying the simulated flowlines. Blue colorscale represents DNA accumulation for experiment and simulation, while grey color scale shows the relative vapor concentration in the simulation. Arrows (right panel) proportionally show the evaporation speed along the interface. **(c)** The simulated and experimentally measured distribution of flow velocities of dissolved beads plotted in a histogram, showing a similar profile. Color scale is equal to (a). **(d)** The maximum relative concentration of DNA increased within an hour to ≈ 30 X the initial concentration, with the trend following the simulation. Error bars are the standard deviation from four independent measurements.

**Figure 3.**
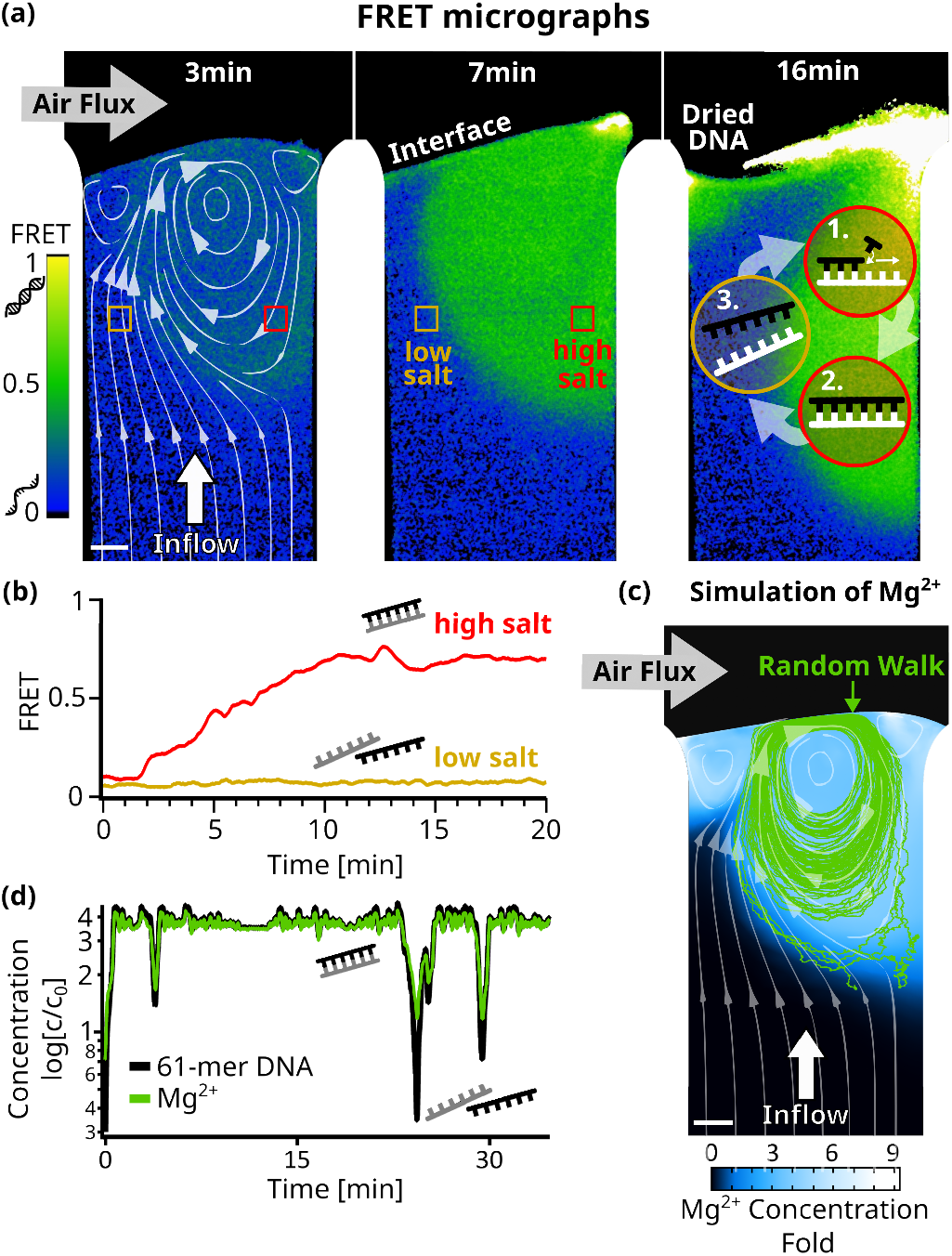
Strand separation by salt cycling. Fluorescence resonance energy transfer measurements revealed cycles of strand separation. **(a)** Micrographs of 24bp DNA FRET pair in the chamber at 45°C. 1 μl sample (5 μM DNA, 10 mM TRIS pH7, 50 μM MgCl2, 3.9 mM NaCl) was subjected to a 3 nl/s diluting upflow of pure water and a gas flow of 230 ml/min across. The induced vortex, shown by the simulated flow lines (left panel), overlays with regions of high FRET indicative of double-stranded DNA. The vortex flow was expected to enable replication reactions by (**1+2**) strand replication in the high salt region and (**3**) strand separation of template and replicate in the low salt region. Fluctuations in interface position can dry and redis-solve DNA repeatedly (see “Dried DNA” in right panel). **(b)** FRET signals confirmed strand separation in low salt regions and strand annealing in high salt regions in (a). After about 10 minutes, DNA and salt accumulated at the interface forming stable and clearly separated regions of low – where the influx from below reaches the interface – and high – located at the vortex – FRET signals. **(c)** Comsol simulation of Mg^2+^ ions (D = 705 μ*m*^2^ /*s* in the chamber agreed with the FRET signal and showed up to 9-fold salt accumulation at the interface. The path of a 61mer DNA molecule from a random walk model is shown by the green lines and the white flowlines are taken from the simulation. **(d)** Concentrations along the DNA molecule path in (c) show oscillations relative to the initial concentration of up to 3-fold for Mg^2+^ and 4-fold for 61mer DNA. This could enable replication cycles, as the vortex provides high salt concentrations for replication, while drops in salt and template concentrations regularly trigger strand separation.

**Figure 4.**
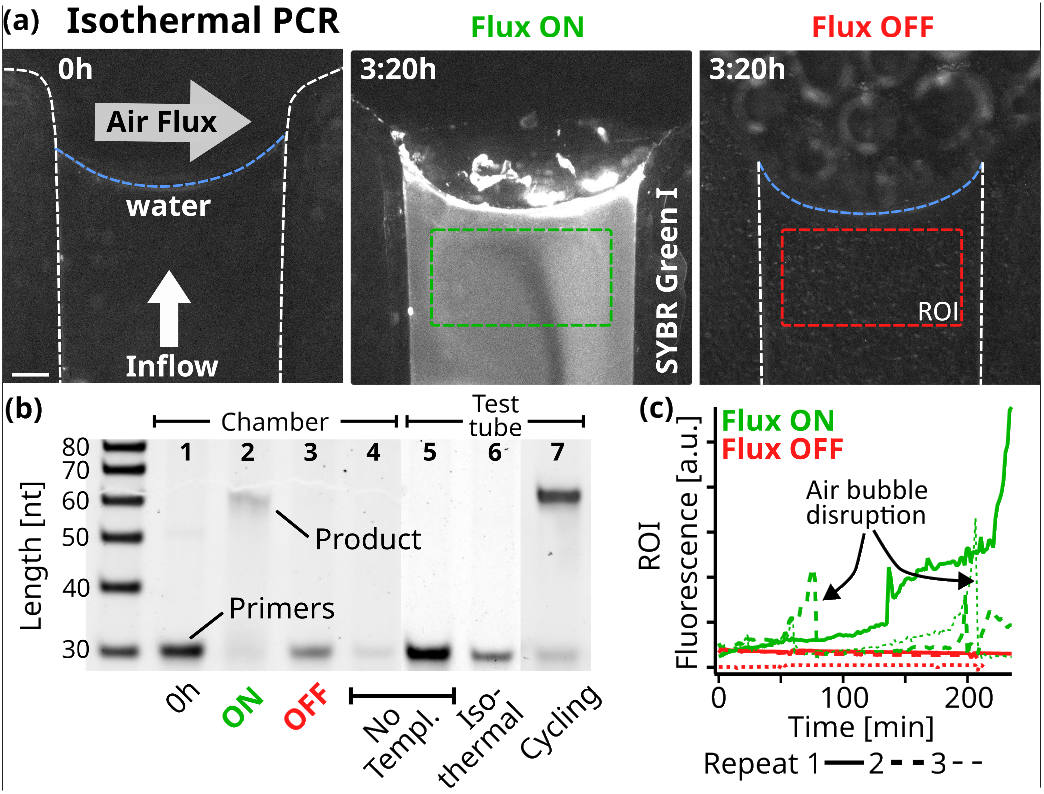
Replication. **(a)** Fluorescence micrographs of the PCR reaction in the chamber. At isothermal 68°C, 10 μl of reaction sample was subjected to a constant 5 nl/s pure water flow towards the interface where a 250ml/min gas flowed perpendicularly. The initial state on the left shows the background fluorescence. Fluorescence increased under flux (middle, after 3:20h), while without flux the fluorescence signal remained minimal (right). The reaction sample consisted of 0.25 μM primers, 5 nM template, 200 μM dNTPs, 0.5 X PCR buffer, 2.5 U Taq polymerase, 2 X SYBR Green I. Scale bar is 250 μm. **(b)** 15% Polyacrylamide Gel Electrophoresis of the reactions and neg. controls. After 4 hours in the reaction chamber with air- and water-flux ON, the 61mer product was formed under primer consumption **(2)**, unlike in the equivalent experiment with the fluxes turned OFF **(3)**. At the beginning of the experiment **(1)** or in the absence of template **(4)**, no replicated DNA was detected. The reaction mixture was tested by thermal cycling in a test tube **(5-7)**. As expected, replicated DNA was detected only with the addition of template: **(7)** shows the sample after 11 replication cycles. The sample was also incubated for 4 hours at the chamber temperature (68°C) yielding no product **(6)**. Primer band intensity variations are caused by material loss during extraction from the microfluidic chamber. **(c)** SYBR Green I fluorescence increased when gas and water flow were turned on, but remained at background levels without flow. Fluorescence was averaged over time from the green and red regions of interest shown in (a). Dotted lines show the data from independent repeats. Air bubbles formed through degassing can momentarily disrupt the reaction. SYBR Green I fluorescence indicates replication, as formed products are able to hybridize.

## Results and Discussion

### Molecule Accumulation at the Gas-Water Interface

We started off by constructing a laboratory model of the rock pore shown in Fig. 1. Here, we focused on the key properties of the system: An upward water flux evaporating at the intersection with the perpendicular gas flux. This leads to an accumulation of dissolved molecules at the interface since they cannot evaporate. Simultaneously, the momentum transfer of the gas flux induces circular currents in water, forcing molecules back into the bulk.

In the following, we analyse how these two effects act on dissolved nucleic acids. For simplicity, we used ambient air as the gas source, enabling us to focus solely on evaporation and the resulting currents. The velocity of the water flowing in was controlled by a syringe pump and chosen to match the velocity of the water evaporating in the given geometry. This ensured reliable and stable conditions in long lasting experiments. For a real early Earth environment we envision a system that self-regulates the water column’s inflow by automatically balancing evaporation with capillary flows. The interface adjusts its position relative to the gas flux, moving closer if the inflow is less than the evaporation rate, or receding if it exceeds it. When the interface nears the gas flux, evaporation accelerates, while moving it away slows evaporation. This dynamic process stabilizes the system, with surface tension ultimately fixing the interface’s position.

Fixed volumes of sample solutions (containing beads, labelled molecules, salts etc.) were always loaded ahead of an influx of pure water, simulating a continuous dilution scenario. The micro scale gas-water evaporation interface consisted of a 1.5 mm wide and 250 μm thick channel that carried an upward pure water flow of 4 nl/s ≈ 10 μm/s. Over this channel, a perpendicular air flow of about 250 ml/min ≈ 10 m/s (Suppl. IV) is guided across. The temperature of the chamber was controlled by a water bath at 45 °C, while a self-built fluorescence microscope provided imaging (See Suppl. Fig. III.1). Two-dimensional finite element simulations were performed to model the diffusion of molecules in water, as well as the flow of water and gas.

First, using a particle tracking algorithm (See Suppl. Sec. V), we measured the flow velocities of individual fluorescent 0.5 μm beads to monitor the dynamics of the water flow as the air-flux streamed across the interface (Fig. 2(a)). As expected, these velocities were dependent on their distance from the channel walls (see Suppl. Fig. VI.1(e)). The beads that were far from the interface were moving at water inflow velocities of 15 μm/s. Closer to the interface, the velocities increased to about 1 mm/s due to the momentum transfer of the gas flow (Supplementary Movie 1). This resulted in a circular flow pattern with the vortex center right below the interface. The flow lines in Fig. 2a) show how the upwards water flux reaches the interface on one side of the vortex, whereas on the opposite side, the beads are pushed back down into the bulk.

The extracted traces in Fig. 2(a) were compared with a finite element simulation. In a two-dimensional projection of the experimental geometry, we modeled laminar gas and water flow, diffusive nucleic acid mass transport in water, interfacial evaporation dynamics, and momentum transfer of gas flowing over the water surface (Suppl. Sec. VI). In agreement with the experimental results, the simulation showed a chamber-averaged water evaporation speed of 10.5 μm/s. The tangential velocity at the interface reached 0.9 mm/s. The modeled flow speed distributions agreed well with distribution of the experimental bead velocities, as shown in Fig. 2(a,c).

To further test our understanding of the dynamics of the system, we imaged fluorescently labeled nucleic acids. The expectation was that the continuous evaporation would lead to an accumulation of the strands at the interface, while the gas flow would induce a vortex analogous to the beads. Both are found to be present in the experiment and agree qualitatively with the finite element model. In our experiment, 2 μl of 5 μM FAM-labeled 63mer DNA were introduced into the system, followed by a continuous diluting pure water inflow. Temperature, water flow and air flow were unchanged from the previous experiment.

Water continuously evaporated at the interface, but nucleic acids remained in the aqueous phase accumulating near the interface. They could only escape downward either by diffusion or by the vortex induced by the gas flowing across the interface, pushing the molecules back deeper into the bulk (See the flow lines in Fig. 2(b) taken from the simulation). As the gas flow continuously removed excess vapor, the evaporation rate remained constant. Thus, except for fluctuations, a stable interface shape should be expected. However, due to the high surface tension of water, the interface is very flexible. As the inflow and evaporation work to balance each other, the shape of the interface adjusts, likely in response to small fluctuations in gas pressure and spatial variations in water surface tension. This is leading to alterations in the circular flow fields below (Supplementary Movie 2).

As these fluctuations are difficult to simulate, we decided to stick with one interface shape, matching evaporation and inflow speeds. The evaporation rate at the interface was therefore set to be proportional to the vapor concentration gradient and varied spatially along the interface between 5 and 10.5 μm/s (See Suppl. Fig. VI.1d)). Using the known diffusion coefficient of 95 μm^2^/s for the 63mer[9], the simulation closely matched the experimental results. In both cases, DNA accumulated in regions with circular flow patterns driven by the gas flux (Fig. 2(b), right panel).

5 minutes after starting the experiment, the maximum DNA accumulation was 3-fold, while after one hour of evaporation, around 30-fold accumulation was observed. Due to molecules residing in very shallow volumes when directly at the interface, the fluorescence signal can vary drastically compared to measurements deeper in the bulk. This can be seen in the fluctuations between independent measurements (See Supplementary Movies 2b,2b,2c), especially around 0.5 h shown in Figure 2(d). The simulated maximum accumulation followed the experimental results and starts saturating after about one hour (Fig. 2(d)).

### Strand Separation Dynamics

As discussed earlier, strand separation is essential for the replication of nucleic acids. Only then can replication become exponential and compete with naturally exponential degradation kinetics. Usually, an elevation of temperature can separate strands but is accompanied with a higher risk for hydrolysis. The chosen isothermal setting requires changes in salt concentration for this process. More specifically, the circular fluid flow at the interface provided by the gas flux, together with Brownian motion, was expected to drive cyclic strand separation by forcing nucleic acid strands through areas of varying salt concentrations.

We used Förster resonance energy transfer (FRET) microscopy to optically measure the strand separation of DNA (Fig. 3). A high FRET signal indicates that two DNA strands are bound, while a low FRET signal indicates that the strands are separated. In this way, FRET becomes an indirect measure of the salt concentration, since a low salt concentration will induce strand separation due to the reduced ionic shielding of the charged DNA or RNA backbones. Specifically, we chose a complementary 24mer DNA pair, with the FRET-pair fluorophores positioned centrally on opposite strands. 1 μL Sample (10 mM TRIS at pH 7, 50 μM MgCl_2_, 3.9 mM NaCl, and 5 μM of each DNA strand) was injected into the chamber and flushed towards the interface by pure water with all other conditions equal to before.

Fig. 3(a) shows micrographs of the recorded FRET values for each pixel (Supplementary Movie 3). Initially, the FRET signal increased near the interface (green), indicating areas where DNA is forming double stranded DNA. This area is localized around the vortex created by the gas flow across the interface. In the upward flow to the left of the vortex, DNA was found to be single-stranded (blue). During the course of the experiment, the low and high FRET regions remained stably separated (Fig. 3(b)). This configuration suggests that the vortex could drive a cycle of replication and strand separation (see the scheme in Fig. 3(a) - right panel).

To confirm this, we simulated the accumulation of Mg^2+^ ions in the chamber (Suppl. VII), since divalent ions have a large effect on the melting temperature of nucleic acids [18]. We then used a Monte Carlo random walk model (Suppl. VIII) to simulate individual 61mer DNA molecules following the vortex and undergoing Brownian motion. Such a path is shown in Fig. 3(c), plotted over the simulated steady-state concentration of Mg^2+^ along with the simulated flow lines. Starting in a region of low Mg^2+^ concentration, the strand enters the vortex created by the gas flow. We have plotted the Mg^2+^ concentration along its path, showing significant salt oscillations of up to 3X the initial salt concentration, capable of inducing strand separation (Fig. 3(d) and Supplementary Fig. VII.2). Rayleigh-Bénard convection cells generate similar patterns to those seen in Fig. 3(c). The oscillations in salt concentration resemble the temperature fluctuations observed in convection-based PCR reactions from earlier studies [32, 33], which also showed that chaotic temperature variations like the salt variations in our system, even enhanced the efficiency of the PCR reaction, compared to periodic ones.

In the experimental conditions used here, RNA would also not readily degrade, even if the strand enters the high salt regimes (See Suppl. Sec. IX). Using literature values for hydrolysis rates under the deployed conditions, we estimate dissolved RNA to have a half life of around 83 days.

When plotting the simulated steady-state concentration of other dissolved – complementary – 61mer DNA molecules along its path, we observed even stronger oscillations of up to 4X the initial concentration. Together with significant drops in Mg^2+^ concentration, this suggests the possibility of exponential replication by strand separation cycles.

### Isothermal Replication with PCR

We saw that nucleic acids and salts accumulated near the interface, but far from the interface, in the bulk below, the concentrations remained vanishingly low due to the diluting inflow of pure water. The air flux induced an accumulation pattern of vortices in which molecules were trapped. The salt and DNA concentration changed cyclically, resulting in periodic strand separation of nucleic acids. Motivated by the above results, we used a model system to test whether nucleic acid replication could actually be implemented in this environment.

We chose to use Taq DNA Polymerase because it does not have a protein-based strand separating mechanism. Starting with a 51mer template and two 30mer primer strands, each with a 5’-AAAAA overhang for detection, the reaction is expected to form a 61mer replicate (Suppl. Sec. X), the same length as the DNA used in the random walk model in Fig. 3(c)&(d). In contrast to standard PCR, which uses thermal cycling to separate the strands, we operated the experiment at isothermal conditions (68°C) and used 10 μl of the reaction mix (0.25 μM primers, 5 nM template, 200 μM dNTPs, 0.5 X PCR buffer, 2.5 U Taq polymerase, 2 X SYBR Green I). This reaction mixture was then exposed to a constant pure water influx of 5 nl/s towards the gas-water interface, matching the rate of evaporation at the interface.

Through the oscillations in salts and DNA observed along the random walk, we expected the 61mer product strand to be able to separate from its respective template strand, enabling exponential replication. The progress was monitored using the intercalating dye SYBR Green I, which binds preferentially to double-stranded DNA [34]. Fig. 4(a) shows fluorescence micrographs of the reaction in the chamber. Initially, minimal fluorescence is seen, indicating that the replicated templates are below the detection limit of SYBR Green. Fig. 4(c) shows how the SYBR Green fluorescence increased after two hours in the displayed region of interest (ROI), recording the increase of replicated DNA forming duplex structures. In other repetitions of the reaction, this increase was sometimes even observed earlier, around the one-hour mark (dotted lines). However, air bubbles nucleated by degassing events, rise and temporarily dry out the channel, interrupting the reaction until the liquid refills the channel (Supplementary Movies 4,4b,4c&5). Despite our best efforts, we were unable to fully prevent this, especially given the high temperatures required for Taq polymerase activity. In an identical setting when the gas- and water flux were switched off, no fluorescence increase was found (See Fig. 4(c) red lines). Fluorescence variations are additionally caused by fluctuations in the position of the gas-water interface, as discussed earlier.

Replication was confirmed under flux with the 61mer product being visible in gel electrophoresis with depleted primers (Fig. 4(b)). With both gas flow and water influx turned off, no product band was found. We verified the replication reaction by repeating the experiments without the addition of the template, primer or DNA in the chamber as well as in a test tube. Suppl. Fig. X.2 shows all independent repeats of the corresponding experiments. No product was detected in any of these cases, ruling out reaction limitations such as primer dimer formation. Primer dimers would form even in the absence of a template strand and would be identifiable through gel electrophoresis. As Taq polymerase requires a significant overlap between the two dimers to bind, this would result in a shorter product compared to the 61mer used here. We also compared the chamber experiment with a regular, temperature cycling based, PCR reaction in a test tube, revealing that in the chamber, about 10-11 cycles of PCR were finished after the 4 hours of experiment (Suppl. X). The findings above confirm that the gas flow at the simulated rock opening was necessary for nucleic acid replication.

### Conclusion

In this work we investigated a prebiotically plausible and abundant geological environment to support the replication of nucleic acids. We considered an isothermal setting of gas flowing over an open rock pore filled with water. Previously, thermal gradients have been used to separate the strands of nucleic acids, risking their degradation. Now, the combined gas and water flow at an open pore trigger salt oscillations. We found that this condition supports oligonucleotide replication. We began by probing the system with fluorescent bead and DNA measurements, finding our results to agree with fluid dynamics theory using finite elements simulations.

While DNA accumulates at the vortex close to the interface, oscillations in nucleic acid and salt concentration are created by a combination of molecular accumulation and interfacial flow, periodically separating nucleic acid strands under chemically gentle conditions. Due to the limitations of RNA-based replication, we probed the environment with protein-driven DNA replication and found isothermal replication in this common geological micro-environment, showing that it provides a setting for early nucleic acid replication chemistry.

Prebiotic chemical reactions such as polymerization of imidazole [20] or 2^′^,3^′^-cyclic phosphate-activated nucleotides [28, 35] will benefit from the reduced RNA hydrolysis in the gentle isothermal replication environment. Most importantly, the combination of actively generated high concentrations by evaporation, dry-wet cycles at the interface caused by interface fluctuations, shielding from UV damage, and the possibility of constant feeding by water influx makes the environment a compelling candidate for implementing the geophysical boundary condition of the early RNA world stage of emergent life.

Furthermore we expect that other gases, such as CO_2_, could establish chemical gradients in this environment. Such gradients have been observed in thermal gradients before [23] and finding similar behaviour in an isothermal environment would be a significant discovery. Physical non-equilibria, such as steep temperature gradients, pose many boundary conditions, decreasing the likelihood of readily finding such a setting on early Earth. This isothermal environment, however, greatly extends the repertoire of prebiotic settings that enable replication on early planets.

## Methods

A microfluidic chamber was created between two sapphire plates (0.5mm thick on the front and 1mm thick on the back), sandwiching a 0.25mm thin Teflon sheet that defined the geometry created by a computer-controlled cutter. The plates were held together by a steel frame bolted to an aluminum back to ensure gas tightness. The back was connected to a water bath (Julabo) to control the temperature. Samples were injected into the chamber using syringe pumps (Nemesys) with tubings inserted into holes in the back sapphire. Gas flow was generated under pressure control using the AF1 dual pump system (Elveflow). Temperature was measured during the experiments with a thermal sensor attached to the back sapphire. A more detailed schematic of the microfluidic chamber can be found in Suppl. IV.

DNA fluorescence measurements were performed in a self-built tilted epi-fluorescence microscope setup using two M490L4 and M625L3 light-emitting diodes (Thorlabs), a 470/622 H dual-band excitation filter (AHF), a 497/655 H dual-band dichroic mirror, and a 537/694 H dual-band emission filter. A more detailed schematic of the setup can be found in Suppl. III. DNA strands were ordered from biomers.net including purification by high-performance liquid chromatography (Suppl. II.1). The strands used for fluorescence quantification of accumulation are (5’-3’)-24bp DNA: *CY5*CGTAGTAAATATCTAGCTAAAGTG, 63bp DNA: *FAM*CCAGCCTCCAGTGCCTCGTATCATTGTG-CCAAAAGGCACAATGATACGAGGCACTGGAGGCTG diluted to 5 μM in Nuclease free water. Images were captured using a Stingray F145B camera (Allied Vision). Bead experiments used 0.5 μM fluorescent microspheres (Invitrogen) diluted 1/2000 in water (Suppl. Sec. V).

2D finite element simulations were performed using COMSOL Multi-physics 5.4. Fluid dynamics were simulated by solving the Navier-Stokes equation in two dimensions. Parameters used are available in table VI.1 in the supplementary Information. The complete description of the model can be found in Suppl. VI.

FRET imaging was performed using a second custom-built fluorescence microscopy setup consisting of light-emitting diodes (M470L2, M590L2; Thorlabs) combined by a dichroic mirror on the excitation side, while an Optoplit II with a ratiometric filter set (DC 600LP, BP536/40, BP 630/50) and a Stingray-F145B ASG camera (Allied Vision Technologies) through a 1X objective (AC254 100-A-ML Achromatic Doublet; Thorlabs) detected and superimposed both fluorescence emission channels (Suppl. III). The DNA sequences used for FRET experiments were: strand 1 5’-CGTAGTAAATAT*FAM*CTAGCTAAAGTG-3’, strand 2 5’-CACTTTAGCTAGAT*ROX*ATTTACTACG-3’. The two labeled complementary strands were diluted from stock solution (100 μM in nuclease-free water) and mixed together to a final concentration of 5 μM in buffer (10 mM TRIS, 50 μM MgCl_2_, 3.9 mM NaCl, pH7). To promote annealing of the two complementary strands, the solution was heated and slowly cooled from 80°C to 4°C (ramp rate of -1°C per 5 s) in a standard thermocycler (Bio-Rad CFX96 Real-Time System) prior to each experiment.

Polymerase chain reaction (PCR) was performed using an AllTaq PCR Core Kit (QUIAGEN). Samples were mixed with 0.5 X AllTaq PCR Buffer, 5 nM template strand, 0.25 μM primers, 200 μM of each dNTP, 2 X SYBR Green I and AllTaq polymerase at 2.5 U/reaction. The reaction in the thermocycler was performed using a temperature protocol of 95°C for 2 minutes for heat activation of the enzyme, then annealing the primers to 52°C for 10 seconds, then 68°C for 10 seconds, and finally 10 seconds at 95°C. This cycle was repeated 40 times (See Suppl. Fig. X.2b)). The reaction in the chamber was performed with 10 μl of the above mixture at 68°C. The solution was also heat activated at 95°C for 2 min followed by an annealing step to 52°C before loading into the chamber. The DNA sequences for the reaction were as follows Template (5’-3’)-51bp DNA: TTAGCAGAGCGAGGTATGTAG-GCGGGACGCTCAGTGGAACGAAAACTCACG, Reverse primer (5’-3’)-30bp DNA: AAAAACGTGAGTTTTCGTTCCACTGAGCGT, forward primer (5’-3’)-30bp DNA: AAAAATTAGCAGAGCGAGGTATGTAG-GCGG.

For PAGE and gel imaging, a 15% denaturing (50% urea) polyacrylamide gel with an acrylamide:bis ratio of 29:1 was solidified with TEMED (tetramethylethylenediamine) and ammonium persulfate. 2 μl of sample was mixed with 7 μl of 2X loading buffer (Orange G, formamide, EDTA), of which 5 μl were loaded onto the gel. Staining was performed with 2X SYBR Gold in 1X TBE buffer for 5 minutes and the gel was imaged using the ChemiDOC MP imaging station (Bio-Rad).

## Data availability

Supplementary data beyond the supplementary material will be given upon request.

## Author contributions

Project conception: P.S., D.B., Research design: P.S., D.B., Methodology development: P.S., Experiments: P.S., E.E. Data analysis: P.S., Programming: P.S., Manuscript writing: P.S., Manuscript reviewing: P.S., D.B., Supervision: D.B. Funding acquisition: D.B.

## Additional information

The online version contains supplementary material available at https://doi.org/10.7554/eLife.100152.1

Funding and acknowledgments

We would like to acknowledge the following agencies for funding: Deutsche Forschungsgemeinschaft (DFG, German Research Foundation) – Project-ID 364653263 – CRC 235, Deutsche Forschungsgemeinschaft (DFG, German Research Foundation) – Project-ID 521256690 – CRC 392, Deutsche Forschungsgemeinschaft (DFG, German Research Foundation) – Project-ID 201269156 – SFB 1032, Volkswagen Initiative’ Life? – A Fresh Scientific Approach to the Basic Principles of Life’, HFSP RGP003/2023, Germany’s Excellence Strategy EXC-2094-390783311, Simons Foundation #327125, European Research Council EvoTrap #787356, ERC-2017-ADG. This work was supported by the Center for Nanoscience Munich (CeNS).

