## Supplementary Information for "Prebiotic Gas Flow Environment Enables Isothermal Nucleic Acid Replication"

Philipp Schwintek[1], Emre Eren[1], Christof Mast[1] and Dieter Braun[1,†]

[1]*Systems Biophysics, Physics department, Center for NanoScience, Ludwig-Maximilians-Universität München, Amalienstraße 54, 80799 Munich, Germany*

#### Contents

I. Movies 2

II. DNA Strands 3

III. Experimental Setups 4

IV. Microfluidic Chamber 5

V. Bead Measurements 6

VI. Finite Elements Simulations 7

VII. Förster Resonance Energy Transfer (FRET) 10

VIII. Random Walk Model 12

IX. Hydrolysis Estimation 13

X. PCR using Taq Polymerase 13

---

†

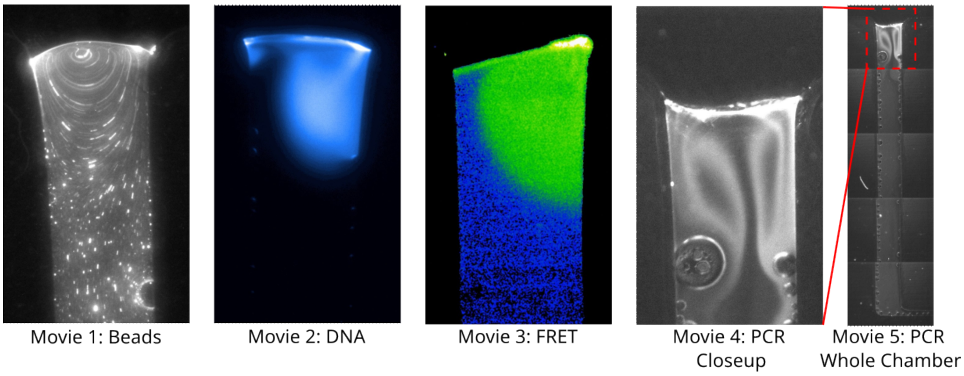


Figure I.1. Snapshots of each supplementary movie.

### Supplementary information I: Movies

The recorded fluorescence images movies which are the basis of the data shown in Fig.2(a), Fig.2(b), Fig.3(a) and Fig.4(a) are provided as supplementary movies.

**Supplementary Movie 1**
Fluorescence beads are used to track the fluid flow shown in Fig.2(a).

**Supplementary Movie 2**
Concentration of fluorescently labelled 63mer DNA is imaged to infer the accumulation at the interface in Fig.2(b). Color scale is the same as in Fig.2(b).

**Supplementary Movie 3**
FRET imaging of dual-labaled DNA strands discriminate between for single stranded DNA in blue and double stranded DNA in green to yellow as detailed in Fig.3(a).

**Supplementary Movie 4**
Top fraction of the chamber used for the figures. SYBR green fluorescence shows the amount of DNA generated in the replication reaction, indicating by a raise of fluorescence how DNA becomes copied for Fig.4(a). In addition, the accumulation at the interface is also seen.

**Supplementary Movie 5**
Whole length of the chamber. SYBR green fluorescence shows the amount of DNA generated in the replication reaction, indicating by a raise of fluorescence how DNA becomes copied for Fig.4(a).

**Supplementary Movie 2b / 2c / 2d**
Individual repeats of Movie 2. Here, the raw data from the fluorescence measurements is shown to underline the interfacial fluctuations.

**Supplementary Movie 4b / 4c**
Individual repeats of Movie 4. Note how gas bubbles formed from degassing travel upwards the channel, drying off the reaction until the channel is filled with liquid again.

### Supplementary information II: DNA Strands

| Length | 5’- Sequence -3’ | Label |
|---|---|---|
| 63mer | ccagcctccagtgcctcgtatcattgtgccaaaaggcacaatgatacgaggcactggaggctg | 5′ FAM |
| 24mer FRET strand 1 | CGTAGTAAATA8CTAGCTAAAGTG | 8 = FAM |
| 24mer FRET strand 2 | CACTTTAGCTAGA8ATTTACTACG | 8 = ROX |
| 51mer Template | TTAGCAGAGCGAGGTATGTAGGCGGGACGCTCAGTGGAACGAAAACTCACG | - |
| 30mer forward primer | AAAAA TTA GCA GAG CGA GGT ATG TAG GCG G | - |
| 30mer reverse primer | AAAAA CGT GAG TTT TCG TTC CAC TGA GCG T | - |

Table II.1. DNA sequences as ordered from biomers.net.

#### Supplementary information III: Experimental Setups

All setups used in this paper were designed and assembled from components purchased from several companies. Figure III.1 shows schematics of both setups used. All specifications below are given in nanometers.

Setup A was built to capture fluorescence signals from beads and labeled DNA and was constructed using SM1 lens tubes (Thorlabs), excitation LEDs M490L4 and M625L3-C4 (Thorlabs), excitation filters (470/622H, AHF) and emission filters (497/655H, AHF), and a dual-band dichroic beamsplitter (497/655H, AHF). The camera (Stingray F-145 B/C) was purchased from Allied Vision. An achromatic doublet (AC254-100-A-ML, Thorlabs) was used as the objective.

Setup B was built to capture FRET. A Zeiss Axiotech Vario microscope body was used and equipped with excitation LEDs M590L4 (yellow) and M470L2 (blue) (Thorlabs) with excitation filters BP588/20 and BP482/29 (Thorlabs) coupled into the same beamline via a DC R 475/40 beamsplitter. A dual-band dichroic mirror (505/606 T) and an Optosplit II with a ratiometric filter set (DC600 LP, BP630/50 and BP 536/40) were coupled to the system to capture the FRET signal of the dyes ROX and FAM. Camera and objective are identical to setup A.

For both setups, fluorescence measurements were performed under curtains to ensure low background noise. Temperature was controlled by a JULABO Corio CD water bath connected to the microfluidic chamber (Suppl. IV) by thermally insulated tubing. The temperature was measured directly at the back sapphire surface of the chamber. Gas flow was generated using an AF1 Dual Pump (Elveflow) system with ambient air as the gas source, and gas flow rate was measured with a flow sensor (FS2000, from IDT). Syringes were driven by a Cetoni syringe pump (Nemesys).

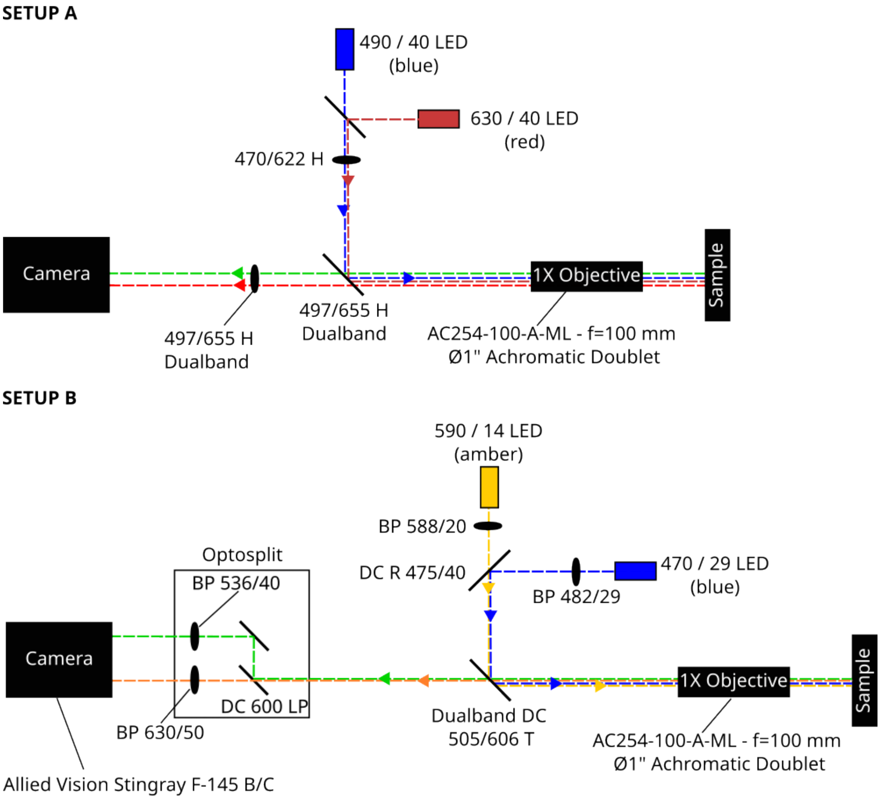


Figure III.1. Sketch of setup A and B used for all fluorescence imaging. Setup A was used for fluorescence measurements containing only fluorescent beads, DNA with a FAM/Cy5 label, and for those containing SYBR Green I. Setup B was used for FRET measurements using the FAM/ROX FRET pair.

### Supplementary information IV: Microfluidic Chamber

The microfluidic chamber used for the experiments in this work was constructed as follows:

Two 60x22mm sapphire plates (front 1mm thick, back 0.5mm thick) were used to sandwich a 250 µm thick Teflon foil from which the channels were cut using a Graphtec CE6000-40 Plus plotter. We used sapphire plates for their higher thermal conductivity compared to normal glass. The back sapphire had four holes to allow gas and water flow in and out of the chamber. The sandwich was held together by the steel frame and aluminum back, which were screwed together with a torque of 0.2 Nm. The aluminum back was held in place on the water bath fixture by magnets. A thin ($25\mu m$) graphite foil was placed between the sapphire back and the aluminum back to increase the thermal conductivity. Figure IV.1 shows an exploded view of the chamber.

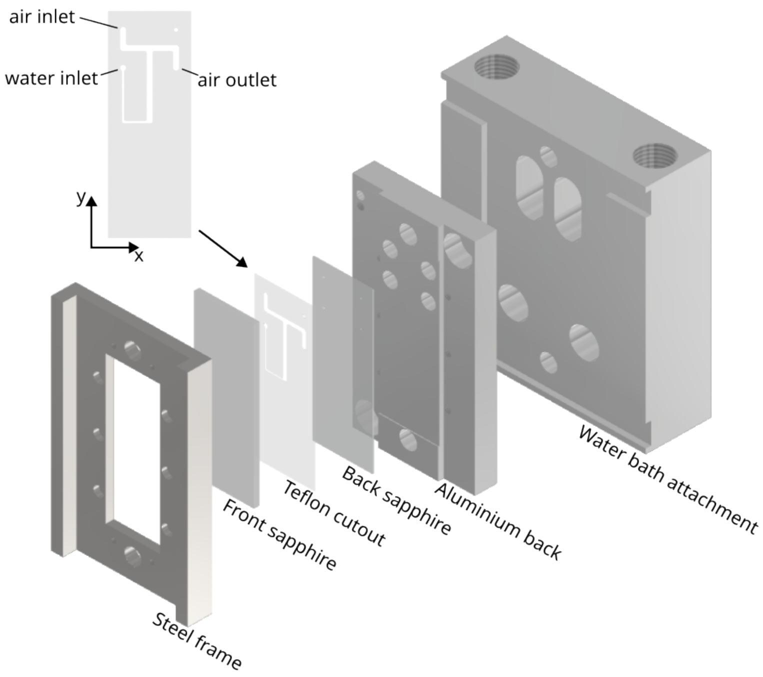


Figure IV.1. Sketch of the microfluidic chamber assembly. Between a steel frame front and an aluminium back attached to the waterbath attachment, a sapphire doublet sandwiches the teflon cutout. The 250µm thick cutout is connected to airflow and waterflow through holes in the back sapphire.

### Supplementary information V: Bead Measurements

A self-written Labview script (Fig. V.1) was used to track the movement of the fluorescent beads. 10µl solution containing 0.5 µm fluorescent beads (Invitrogen, Eugene, Oregon, USA, Lot: 31373W), diluted 1 to 2000, were loaded into the chamber and subjected to a pure water upward flow of 3 nl/s and a perpendicular gas flow of 230ml/min (Supplementary Movie 1). Images were captured using a 50ms exposure time, resulting in approximately 20 fps. Our setup did not allow very fast beads to be traced, as the maximum frame rate of 20fps did not capture particles faster than about 1mm/s. The 2D map of traced velocities therefore has some dark spots near the interface where the fastest beads are located.

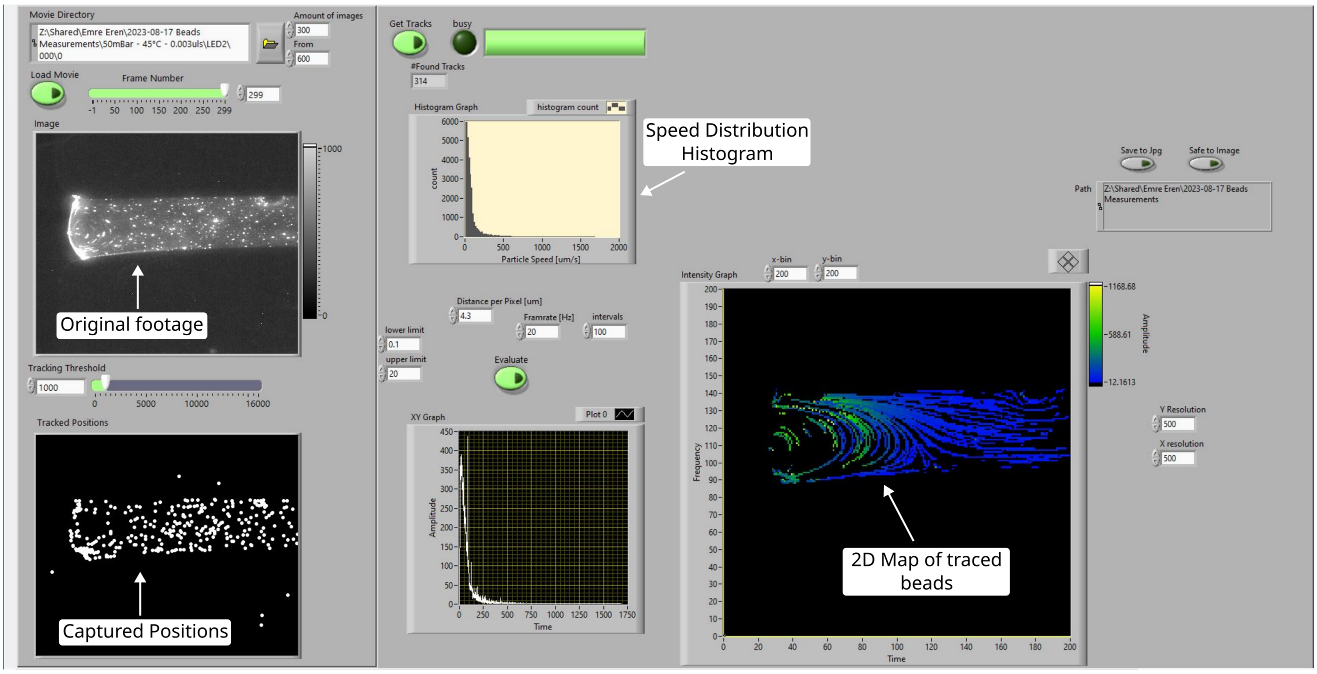


Figure V.1. Screenshot of the user interface of the self-written LabView script used for particle tracking. A series of microscopy images are loaded and beads are located and positional changes between images are traced. Statistics of particle speeds spanning hundreds of images are calculated using imaging framerate and previously measured length/pixel values. A 2D map of the obtained traces converted to the corresponding speed is generated as well.

### Supplementary information VI: Finite Elements Simulations

Finite element simulations were performed using COMSOL Multiphysics 5.4. A 2D geometry was designed using the same parameters as the Teflon cutout for the experimental chamber. Since the dynamics of the experiment take place mainly in the x-y plane, the system was simulated without the z dimension, focusing on the key properties. The geometry is coupled to a gas inlet and outlet as well as a water inlet from the bottom, analogous to the experiment (Suppl. VI). The normal gas inflow has been set to experimentally measured values (about 236 ml/min), resulting in velocities up to about 12 m/s. The gas outlet uses pressure as its boundary condition, releasing as much gas as necessary to maintain a constant pressure. The system is assumed to be laminar, since the velocities don't exceed Mach ¡ 0.3.The transport of water vapor in the gas is coupled at the top, using the stationary velocity field previously established. Simultaneously, the stationary velocity field of the laminar upward flow of water was calculated and coupled with the time-dependent transport of dilute species, in this case dissolved DNA or salts.

For both the gas and the water, Navier-Stokes-Equations were solved under the assumption that the flows are laminar due to their relatively low speed to viscosity ratio (Reynolds number). The flows can therefore be considered incompressible, the density constant and the continuity equation reduces to the condition:

$$\nabla \cdot \boldsymbol{u} = \boldsymbol{0}\,. \tag{1}$$

with $\rho$ denoting the mass density and $\boldsymbol{u}$ the velocity field. The Navier-Stokes equation then reduces to

$$\rho(\boldsymbol{u} \cdot \nabla)\boldsymbol{u} = \nabla \cdot [-p\boldsymbol{I} + \eta(\nabla\boldsymbol{u} + (\nabla\boldsymbol{u})^T] + \boldsymbol{F} = \boldsymbol{0}\,, \tag{2}$$

with $p$ being pressure, $\boldsymbol{I}$ the unity tensor, $\eta$ the fluid dynamic viscosity and $\boldsymbol{F}$ the external forces applied to the liquid. The reference pressure was set to 1[atm], the reference temperature was 45°C and all surfaces, except the gas-water interface, are described as non-slip boundary conditions. The diffusion dependent transport of diluted species was simulated using Fick's law and convection due to the laminar flow fields:

$$\frac{\partial c_i}{\partial t} + \nabla \cdot (-D_i \nabla c_i + \boldsymbol{u} c_i) = 0 \tag{3}$$

Equivalently, equation 2 and 3 are used for the dynamics of the gas channel. The boundary condition to combine the gas-flow with the water-flow is embedded in the gas-water interface: The velocity field components in x- and y-direction of the gas- as well as water-flow are required to be equal at the gas-water interface:

$$\boldsymbol{u} = \boldsymbol{u2} \tag{4}$$

where $\boldsymbol{u}$ describes the vector field of water, while $\boldsymbol{u2}$ denotes the vector field of the passing gas. The interface acts as a sliding wall, moving in the x-direction of the gas-flux to emulate the momentum transfer of the wind to the water surface. To simulate the evaporation of water into the gas phase, we used the August equation, describing the relation between saturation vapor pressure and temperature:

$$P_{sat} = \exp\left\{20.386 - \frac{5132K}{T}\right\}[mmHg] \tag{5}$$

The saturation concentration of water vapor therefore is

$$c_{sat} = \frac{P_{sat}}{R \cdot T} \tag{6}$$

,where R denotes the ideal gas constant and T the temperature. At the interface, the boundary condition reads:

$$c_{vapor} = c_{sat}, \tag{7}$$

while at the ceiling of the gas-channel, far away from the interface, the concentration is set to:

$$c_{vapor} = h \cdot c_{sat}, \tag{8}$$

where h denotes the relative humidity in percent. Furthermore, the speed of evaporation at the interface is proportional to the vapor concentration gradient:

$$\boldsymbol{v}_{\text{evap}} = -D_{vap} \cdot \frac{M}{\rho} \cdot \nabla c_{vap} \tag{9}$$

where M represents the molar mass of water, $\rho$ the density of water and $D_{vap}$ the diffusion coefficient of vapor.

At the inlet, the vapor concentration in the gas is set to a constant humidity dependent value (See table VI.1 for a detailed parameter list).

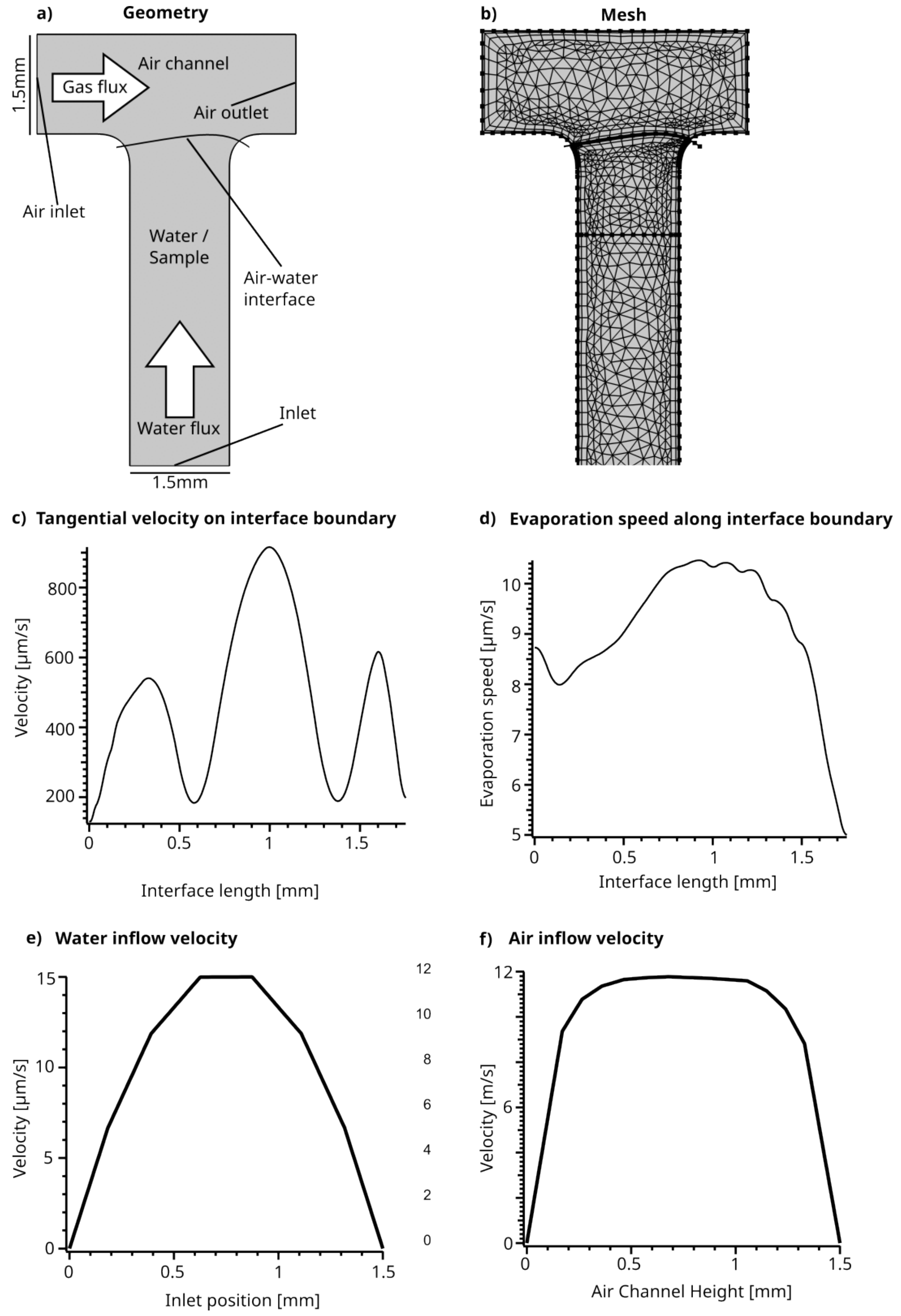


Figure VI.1. **a)** The geometry as it is used for the simulation. **b)** The geometry of the system after it has been meshed.**c)** Tangential velocity directly at the interface. x-Axis goes from the left-most point of the interface (see a)) to the right-most. The velocity is induced by the momentum transfer of the gas brushing across the interface (boundary condition equation 4). **d)** Evaporation speed along the interface. **e)** Parabolic flow profile at the inlet of the chamber in a). **f)** Parabolic flow profile of the gas flow measured at the inlet

| **Parameter** | **Value** | **Description** |
|---|---|---|
| $D_{vapor}$ | (21.2E-6)*(1/1[K])*(1 + (0.0071*(T - 273))) $[m^2/s]$ | Diffusion Vapor |
| $D_{63mer}$ | $643*n^{-0.46}[\mu m^2/s] = 95.6\ \mu m^2/s$ | Diffusion Coefficient of a 63mer DNA Strand [9] |
| $D_{Mg^{2+}}$ | $705\ \mu m^2/s$ | Diffusion Coefficient of $Mg^{2+}$ [36] |
| $c(vapor)_0$ | humidity*0.01*(exp(20.386- (5132[K]/T))[mmHg]) / (R * T) | Initial Vapor concetration |
| Humidity | 40 % | Ambient relative humidity |
| $p_{sat}$ | (exp(20.386 - (5132[K]/T))[mmHg]) | Vapor saturation pressure |
| $c_{sat}$ | $p_{sat}$/(R*T) | Vapor saturation concentration |
| $M_{vapor}$ | 0.0180 [kg/mol] | Molar mass of vapor |
| $M_{water}$ | 18.01528[g/mol] | Molar mass of water |
| T | 45°C | Temperature |

Table VI.1. **Parameters used for the finite elements simulation.** Final set of parameters used to simulate the system. Water-specific parameters such as dynamic viscosity or density were taken from inbuilt features of COMSOL Multiphysics 5.4..

### Supplementary information VII: Förster Resonance Energy Transfer (FRET)

To measure the FRET signal in our microscope, we used an alternating illumination protocol (Supplementary Movie 3). The FRET pair FAM-ROX was excited by two LEDs in rapid succession. The blue LED excited the donor dye (FAM), while the acceptor (ROX) can only be excited indirectly while both dyes are in the FRET region. The yellow LED excited only the acceptor dye (ROX). Individual images of each illumination were captured using an Optosplit II to separate the individual emission wavelengths of FAM and ROX before they reached the camera. This allowed the emission of FAM and ROX to be captured simultaneously for each of the two illuminations, providing four images for each time point: DD, DA, AA, and AD (see table VII.1 for details). The spatially averaged, temperature-dependent, crosstalk- and artifact-corrected FRET signal was calculated using the equation 10[9]. Crosstalk between the two channels (aa(T) and dd(T)) was calculated in separate experiments using the same setup parameters with the equation 11 and 12, respectively. The data used for the crosstalk calculations are shown in Fig.VII.1. To test how different salt concentrations affect the FRET signal, we performed melting curves of different salt concentrations (See Supplementary Figure VII.2). We found that the sodium concentration has little effect on the melting temperature, while $Mg^{2+}$ strongly influences the hybridization state. In the FRET experiment in Figure 3, the initial $Mg^{2+}$ concentration was 50 µM at 45 °C. In this state the double stranded fraction is about 0.3. When the salts accumulated at the interface, salt concentrations increased up to 9 fold, strongly changing the double stranded fraction to around 0.8.

| Channel | Excitation | Emission | Label |
|---|---|---|---|
| DD | FAM - 470nm | FAM - 536nm | FAM/ROX |
| DA | FAM - 470nm | ROX - 630nm | FAM/ROX |
| AA | ROX - 590nm | ROX - 630nm | FAM/ROX |
| AD | ROX - 590nm | FAM - 536nm | FAM/ROX |
| $AA_A$ | ROX - 590nm | ROX - 630nm | ROX |
| $DA_A$ | FAM - 470nm | ROX - 630nm | ROX |
| $DD_D$ | FAM - 470nm | FAM - 536nm | FAM |
| $DA_D$ | FAM - 470nm | ROX - 630nm | FAM |

Table VII.1. Channel definitions for FRET calculation. First capital letter denotes the excitation wavelength (D = Donor, A = Acceptor), second the measured emission wavelength and the subscript stands for the label used in a separate experiment to determine crosstalk related artifacts.

$$FRET(T) = \frac{DA(T) - dd(T) \cdot DD(T) - aa(T) \cdot AA(T)}{AA(T)} \tag{10}$$

where dd(T) and aa(T) represent the non-FRET artifacts (crosstalk) in the DA and AA channels and are defined as:

$$dd(T) = \frac{DA_D(T)}{DD_D(T)} \tag{11}$$

and

$$aa(T) = \frac{DA_A(T)}{AA_A(T)} \tag{12}$$

Before each experiment, a melting curve of the FRET strands was performed inside the experimental setup chamber. The melting curve was used to normalize the FRET signal to 0 and 1 using the following equation:

$$FRET_{norm}(T) = \frac{FRET(T) - \alpha}{\beta} \tag{13}$$

where $\alpha = min(FRET(T))$ and $\beta = max(FRET(T))$.

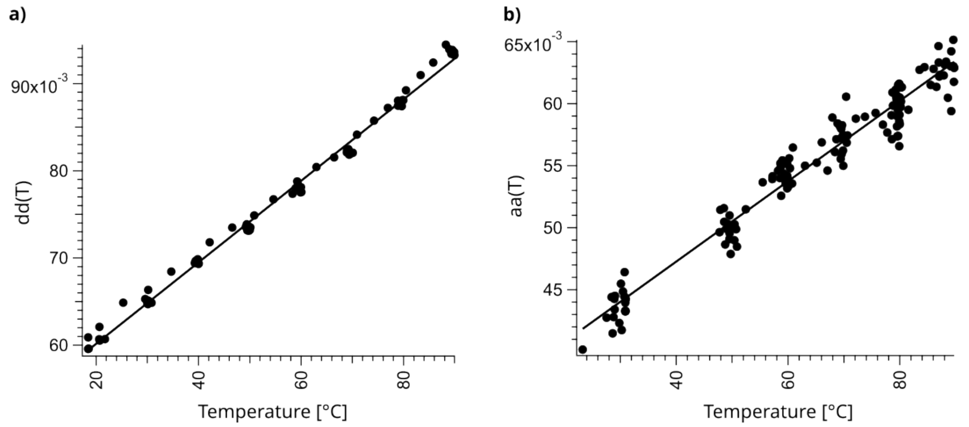


Figure VII.1. Crosstalk between donor and acceptor channel, **a)**: dd(T) and **b)**: aa(T), plotted as a function of temperature and fitted linearly.

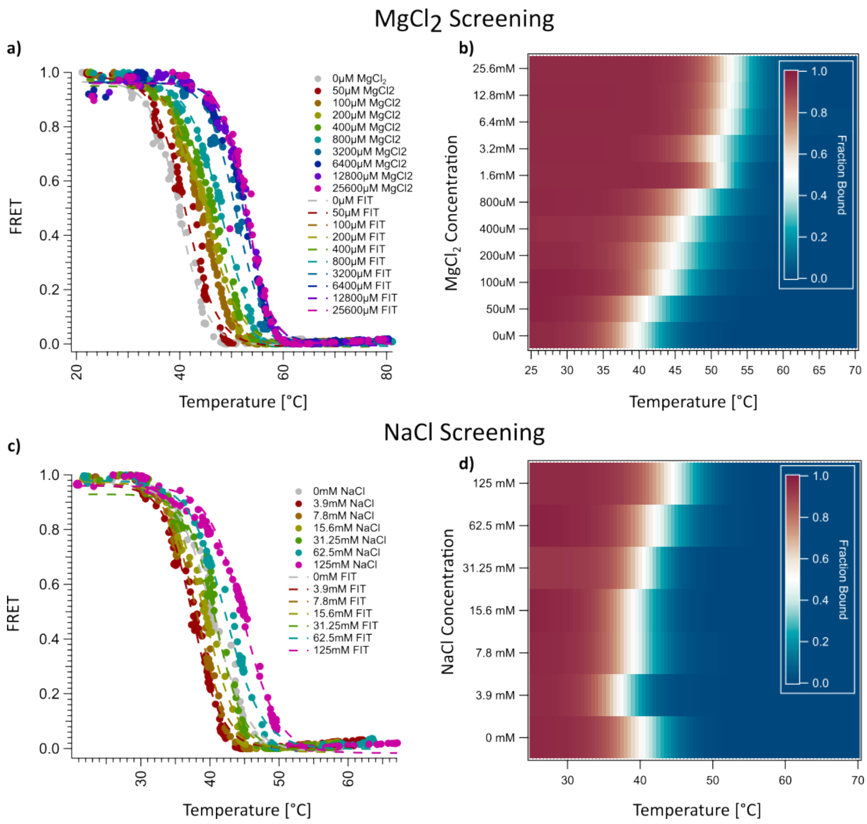


Figure VII.2. Melting curves performed in the FRET setup using the 24mer strands labelled with ROX and FAM respectively. a) Raw data of the melting curves of different $MgCl_2$ concentrations with $\alpha$ and $\beta$ already applied for normalization. b) Data from a displayed as a heatmap. The white areas display a fraction bound of 0.5 corresponding to the melting temperature $T_m$. Note that small oscillations in $Mg^{2+}$ strongly influence the melting temperature, which can enable strand separation at isothermal settings. c) Raw data of the melting curves of different NaCl concentrations with $\alpha$ and $\beta$ already applied for normalization. d) Data from c displayed as a heatmap. The white areas display a fraction bound of 0.5 corresponding to the melting temperature $T_m$.

### Supplementary information VIII: Random Walk Model

For the random walk model, we first used the existing Comsol simulation and ran the simulation with $Mg^{2+}$ ions and a 61mer DNA as the dilute species (the diffusion constant of $705 \frac{\mu m^2}{s}$ for $Mg^{2+}$ at 25°C was taken from [36] and the diffusion constant of $97.04\ \frac{\mu m^2}{s}$ for a 61mer DNA strand from [9]). The simulation was performed in the same chamber, with the same characteristics and settings as in Suppl. Sec. VI. The resulting stationary salt and DNA concentration fields after 2 hours were exported as a 2D table with 200 values in x-direction and 200 values in y-direction, representing the whole simulated geometry. The same was done for the stationary laminar flow field in x-direction and y-direction induced by the air flow across the gas-water interface. Values were linearly interpolated for points between values from the grid.

Then, a self-written LabView script was used to simulate the Brownian motion of a particle with a chosen diffusion constant starting at a random position in the chamber and propagating along the flow vector field in 10ms time steps. To do this, we look at the random square displacement of a particle with diffusion constant D:

$$x^2 = Dt \tag{14}$$

As this particle can move in two directions, left and right this becomes

$$x^2 = 2Dt \tag{15}$$

Expanding this into two dimensions, we get:

$$x_{2D}^2 = x_x^2 + x_y^2 \text{ and therefore: } x_{x,y} = \sqrt{x_x^2 + x_y^2} = \sqrt{2Dt + 2Dt} = \sqrt{4Dt} \tag{16}$$

We then inserted the diffusion constant of a 61mer DNA[9] of $97.04 \cdot 10^{(-12)} \frac{m^2}{s}$ and a random unit vector phi with values between [-1,1]. $\vec{u}$ and $\vec{v}$ represent the exported laminar flow field data from Comsol:

$$\text{x displacements:} \sqrt{4 \cdot 97.04 \cdot 10^{(-12)} \cdot dt} \cdot phi + dt \cdot \vec{u} \tag{17}$$

$$\text{y displacements:} \sqrt{4 \cdot 97.04 \cdot 10^{(-12)} \cdot dt} \cdot phi + dt \cdot \vec{v} \tag{18}$$

The particle was then displaced according to equations 17 and 18 with a timestep of 10ms for a total of 35 minutes. Along its path, the respective local $Mg^{2+}$ and 61mer DNA concentration were plotted, yielding the graph displayed in main text Fig.3(d). The path was overlayed with the original Comsol simulation graphic of the salt concentration distribution in Figure 3(c).

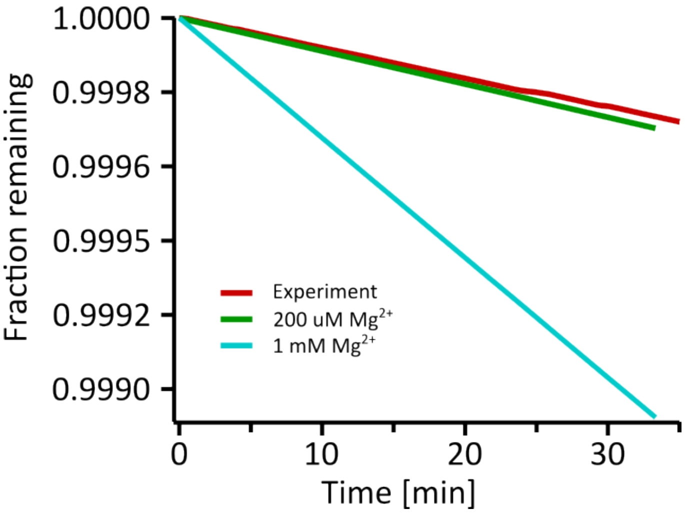


Figure IX.1. Theoretical hydrolysis of RNA in the deployed experimental conditions calculated for the Monte-Carlo trace of Fig. 3 as well as for constant $Mg^{2+}$ concentrations.

#### 130 Supplementary information IX: Hydrolysis Estimation

To estimate the hydrolysis rate of a 24-mer RNA strand deployed in the conditions used in Fig. 3 (45 °C, pH 7, 50 - 200 µM $Mg^{2+}$, 4 - 12 mM NaCl), we used a model from literature to derive the hydrolysis rate in dependence of ion concentrations,
131 temperature and pH [21]:

$$k_{hyd}[1/min] = L \cdot k_{bg} \cdot 10^{0.983\cdot(pH-6)} \cdot 10^{-0.24\cdot(3.16-[K^+])} \cdot 10^{(0.07\cdot(T-23))} \cdot 3.57 \cdot [K^+]^{-0.419} \cdot 69.3 \cdot [Mg^{2+}]^{0.8} \tag{19}$$

Here, $L$ denotes the length of the oligomer (24 in this case), $k_{bg}$ represents the a background hydrolysis rate of 1.3E-9 [1/min], and all concentrations ([$Mg^{2+}$ and [$K^+$]]) are given in Molar. We used $Na^+$ instead of $K^+$ in the experiment, but assumed here,
132 that the $K^+$ correction would be similar for this calculation.

Using the trace of the Monte-Carlo random walk shown in Fig. 3(c), we calculated the hydrolysis rate from equation 19 for
133 each time step of 10 ms iteratively. The resulting relative fraction of remaining oligomers over time is plotted in Fig. IX.1. It
134 also shows two comparison graphs, where a constant $Mg^{2+}$ concentration was assumed over the same course of time. In all cases
135 RNA is very stable. With an average hydrolysis rate of 7.987E-6 1/min during the experiment, RNA is approximated to have a
136 halftime of 2.09E+03 hours. Even at 1 mM $Mg^{2+}$, the halftime reduces to 516 h with a rate of 3.231E-5 1/min. The maximum
137 concentration of $Mg^{2+}$ in the random walk is around 4 X the starting concentration of 50 µM. The static hydrolysis curve at
138 200 µM $Mg^{2+}$ is also shown in Fig. IX.1 with a hydrolysis rate of 8.91703E-6 1/min and a halftime of 1870 h. After 35 min
139 under the experimental conditions of Fig. 3, not more than 1 ‰ of the initial RNA would have hydrolysed. Using a per-base
140 copying rate of a ribozyme polymerase [37, 38] of 72 $h^{-1}$, a 24-mer strand would be copied every 20 minutes. The timescale of
141 replication using prebiotically plausible machinery therefore out-competes the hydrolysis in the experimental settings used here.
Even assuming a base extension rate of 1 $h^{-1}$ for non-enzymatic replication [39], one 24-mer strand would get copied once per
142 day, out-competing the estimated hydrolysis.

Overall the conditions deployed in the experiment are not harsh on RNA. The vortex brings the nucleic acids down to regimes
143 of low salt, further increasing their lifetime repeatedly.

#### 144 Supplementary information X: PCR using Taq Polymerase

Replication reactions were performed using the AllTaq PCR Core Kit (QIAGEN). Each reaction contained 2.5U of AllTaq polymerase, 2X SYBR Green I, 5nM template, 0.25µM of each primer, 200µM of each dNTP and 0.5X PCR buffer (contains
145 Tris HCl, KCl, $NH_4SO_4$ and $MgCl_2$).

To distinguish the template from the product strand on a PAGE image, we have added an overhang of 5A's to the 5' end of each primer. The 51mer template will be extended with the product strand then becoming a 61mer. Figure X.1 shows a scheme of the replication cycle. Steps A to E represent the phase of replication in which the original 51mer template is extended step by step, first to a 56mer and finally to the 61mer product. Once the template is extended, the reduced replication cycle F to G (Fig.X.1 green boxes) begins, in which the concentration of template and product strands increases exponentially. Once the initial amount of template (5nM) is consumed, the reaction can only be represented by the reduced scheme.

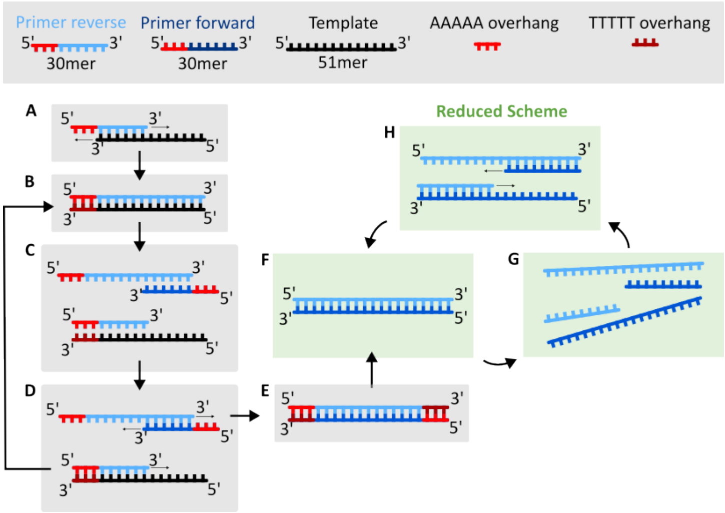


Figure X.1. Complete schematic of the replication reaction using Taq polymerase. **A** The 30mer reverse primer binds to the 51mer template. The 5-A overhang remains unbound. **B** Taq polymerase adds nucleotides from 3’ to 5’ and a complete double strand is formed, now containing 56 base pairs. **C** After strand separation, the newly formed 56mer intermediate product with a 5-A overhang at the 5’ end and the intermediate 56mer template with a 5-T overhang at the 3’ end are bound by the primers. **D** In one case, Taq elongates the forward primer bound to the intermediate product and proceeds to step E. In the other case, elongation of the reverse primer bound to the intermediate template leads back to step B. **E** The result of the extension is a new product and a new template of 61 bases each. From here, the cycle enters the ”reduced scheme”. The newly formed products, together with the original primers, now replicate exponentially: **F** The double strand formed in E can now be considered as product and template for the reduced scheme. **G** After de-hybridization, both primers can anneal to template and product. **H** Taq extends again from 3’ to 5’, forming two new double strands of template and product, doubling their amount and completing one cycle of exponential amplification.

To confirm that the reaction in the chamber followed the predicted scheme, we performed a series of control experiments. Fig.X.2 shows the resulting PAGE gels and the temperature protocol. The experiment was performed either in the microfluidic chamber (Supplementary Movies 4 & 5) or in a test tube inside a thermocycler (Fig.X.2a)). After a heat activation step of 95°C for Taq polymerase, the temperature was kept constant at 68°C in the microfluidic chamber, while in the thermocycler we followed the PCR protocol for Taq, in which the sample underwent multiple cycles of replication (Fig.X.2b)). After 95°C, the primers are given time to anneal by cooling the sample to 52°C, followed by a replication step at 68°C, where Taq is at its peak performance. This temperature protocol is then repeated an additional 39 times to complete the 40 cycles, and the sample is then cooled to 4°C, extracted, and then stored at -20°C until PAGE analysis.

Fig.X.2(c) shows the PAGE results for the test tube samples in the thermocycler. On the left, the triplicate of the full sample (conditions shown in Fig.X.2(a)) shows primer consumption and the formation of a product band in all cases. To be sure that Taq is not forming an unwanted side product, such as primer dimers, we repeated this experiment without adding the template strand, and indeed no product strand can be detected. Furthermore, the reaction cannot proceed to generate product without having both the reverse and forward primers. In the PAGE gel on the right, experiment triplicates are shown without the forward primer, without the reverse primer, or without primers at all. As expected, no product was formed in any of these cases. Without the addition of DNA (no primers and no template), Taq polymerase does not form a new strand. As a further negative control, we kept a complete sample in the chamber at isothermal 68 degrees C, analogous to the experiment, and observed no product formation. This is particularly interesting because it shows that without the microfluidic chamber environment, the replication cycle cannot be completed and the reaction is halted, further emphasizing the need for salt cycling in the chamber experiment.

Fig.X.2(d) shows the PAGE gels of all experiments performed in the microfluidic chamber. While the full sample replicates 2 and 3 show product formation, no product is observed without the addition of the template strand to the reaction mix. When the reaction is run without the reverse primer, forward primer, primers in general, or no DNA at all, no product formation is observed. To show that the replication reaction is only possible when both the gas flow and the water flow are turned on, we repeated the experiment without any fluxes turned on. Here, the full sample was placed in exactly the same microfluidic chamber and kept at isothermal 68 degrees C as in the other chamber experiments. However, without upconcentration at the interface and without continuous stirring by the gas flow, no product formation can be observed.

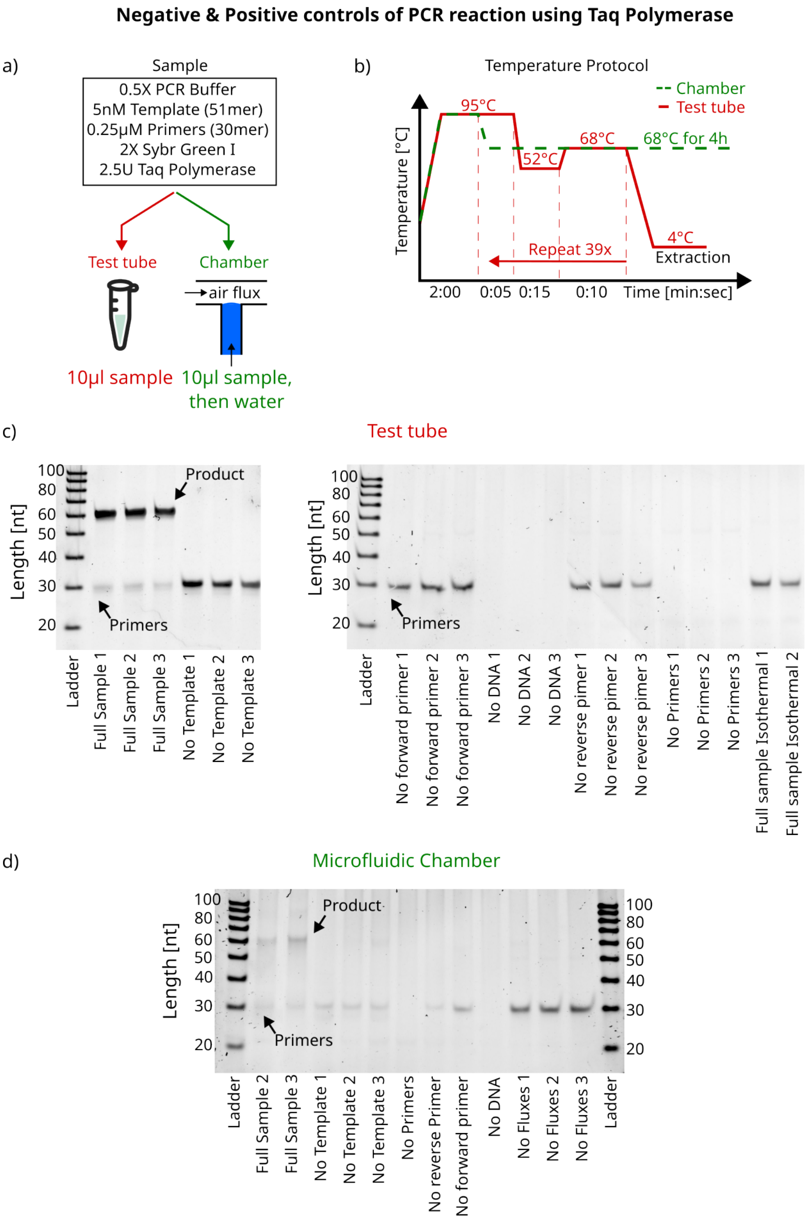


Figure X.2. PAGE analysis of Taq PCR reactions. **a)**Sample composition used for the PCR reaction. The sample is either placed in a test tube (Eppendorf tube) and the temperature is controlled in a thermocycler (red text), or 10 µl of the sample is loaded into the microfluidic chamber prior to dilution water inflow (green text). **b)** Temperature protocol for the two individual experiments. In the thermocycler (red line), the sample undergoes a heat activation step at 95°C, followed by an annealing step at 52°C for 15s, then a replication step at 68°C for 10s. This is repeated 40 times before the sample is extracted and stored at -20°C prior to loading on a gel. Chamber experiments are performed at isothermal 68°C after the same heat activation step and extracted after 4 hours. **c)** PAGE images for the test tube samples. "Full sample isothermal 1&2" samples have the same temperature protocol as the chamber samples. A slight band may be visible around 51nt caused by the 5nM of the 51mer template. The numbers indicate the nth replicate of the experiment. **d)** PAGE image of the chamber control experiments. Primer fluorescence varies between samples, which is caused by variability during sample extraction or primer consumption by Taq (in case of "Full Sample").

Furthermore, we were interested in how many full PCR cycles the sample underwent in the chamber compared to regular PCR using temperature cycling in a test-tube. Therefore we performed the experiment in a test-tube with 10 different amount of temperature cycles as displayd in Figure X.2b). We then compared these results with the samples extracted from the air-flux chamber after 4 hours. Figure X.3a) shows the corresponding 15% PAGE image. However, due to losses during the extraction of the sample from the microfluidic chamber, only comparing the gel band intensity of the product from chamber to test-tube

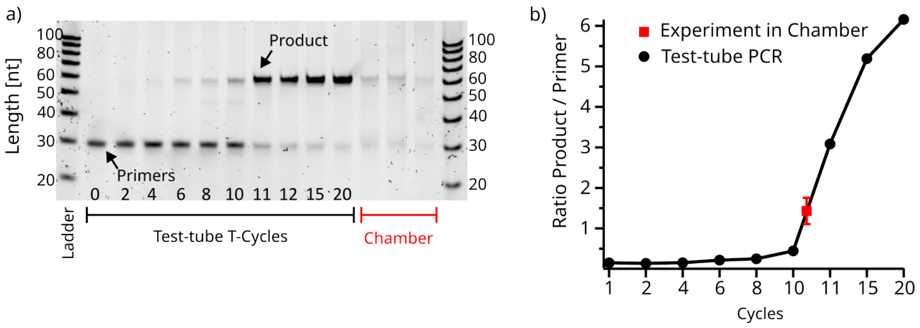


Figure X.3. Comparison of test-tube temperature cycling vs. chamber experiments. **a)** Full Sample (10 µL of 2.5U of AllTaq polymerase, 2X SYBR Green I, 5nM template, 0.25µM of each primer, 200µM of each dNTP and 0.5X PCR buffer) were subjected to various amounts of temperature cycles, as displayed in X.2b). The more cycles are performed, the more primers are consumed to form the product strand. The three experiments performed in the chamber show a generally lower gel intensity, which is due to losses during sample extraction from the microfluidic chamber. **b)** Comparison of Product/Primer intensity-ratio of the test-tube sample to the extracted chamber samples. This reveals that in the 4h chamber experiment about 10-11 cycles were performed. Errorbar of the experiment data point is the standard deviation of the 3 chamber samples in b).

is not representative, because the corresponding primer band intensity does not match any of the test-tube samples. To account for this, we calculated the ratio of product to primer intensity for all lanes (Fig. X.3b)). Since losses during extraction are the same for primer as well as for product strands, the ratio of product to primer strands stays unaffected. This reveals that inside the microfluidic chamber, with air- and water-fluxes turned on, after 4 hours, 10-11 full cycles of replication were performed. Gel intensities were extracted using a self-written LabView tool.
